# Disruption of HIF1Α translational control attenuates the HIF-dependent hypoxic response and solid tumour formation *in vivo*

**DOI:** 10.1101/2022.11.02.514731

**Authors:** Jill E Hunter, Oliver McHugh, Gabrielle B Ecclestone, Fraser Child, Hannah Mearns, Georgia Robson, Molly Dadzie, Sonia Rocha, Neil D Perkins, Niall S Kenneth

## Abstract

Adaptation to reduced oxygen availability is mediated by the hypoxia-inducible factor (HIF) family of transcription factors. The activity and availability of HIF proteins is primarily driven by the stability of the HIF alpha subunits. However, it is becoming increasingly apparent that preferential translation of HIF1α mRNA is also necessary for full activation of the HIF1-dependent hypoxic response. Consequently, the mechanisms controlling HIF1α translation are of equivalent importance to the proline hydroxylase-dependent degradation pathways. Here we investigate the role of the 5’UTR of the HIF1α mRNA in controlling preferential translation of endogenous HIF1α in hypoxic cells. CRISPR/Cas9-mediated genetic deletion of the 5’ UTR of HIF1α results in reduced HIF1α levels following hypoxia, without alteration in mRNA or protein stability. HIF1α mRNA lacking the 5’UTR was efficiently translated in adequately oxygenated cells but this was inhibited during hypoxia, consistent with the global block on protein synthesis. The HIF1α translational defect observed in cells missing the 5’UTR led to reduced viability in hypoxic conditions *in vitro* and an impaired ability to form solid tumours in murine xenografts. Prevention of preferential HIF1α translation limits the duration and intensity of the HIF-dependent hypoxic response and disrupts the formation of solid tumours. Together these results demonstrate the importance of translation control over HIF1α and suggest that strategies to inhibit preferential HIF1α protein translation in hypoxic cancer cells will be an effective strategy to limit the growth of solid hypoxic tumours.

## INTRODUCTION

Hypoxia can occur as a consequence of low atmospheric oxygen or locally in tissues due to inflammation, ischemia, injury, or poor vascularisation, something that particularly evident in solid tumours [1, 2]. As cells encounter hypoxia there is an adaptive switch in energy metabolism coupled with a rapid change in the gene expression program to restore oxygen homeostasis [3]. Hypoxia-dependent changes in translation rates and protein degradation rates are described in many studies but how cells select mRNAs for preferential translation in hypoxic cells is not yet well-understood [4, 5].

Cells can either control mRNA translation on a global level, with the translation of the bulk of the mRNAs being regulated *en masse*, or be mRNA specific, wherein the translation of stress-responsive and adaptative mRNAs are modulated in a more precise manner [6-8]. Since insufficient oxygen in hypoxic cells results in decreased capacity for energy production there is a general suppression of energy expensive processes such as protein synthesis [4, 5, 9]. Hypoxia-dependent repression of protein synthesis occurs mostly at the levels of translation initiation, generally considered to be the rate-limiting step of protein translation [4, 5, 8]. However, while cells are conserving energy by limiting non-essential processes, mRNAs of stress-responsive genes must be actively translated to promote the expression of genes essential to the adaptive response to hypoxia [4, 5, 8]. The cellular signalling pathways that determine this selection, and the defining characteristics of the individual mRNAs that control their translation rates, are not well-understood [4].

One of the best-described mRNAs preferentially translated during hypoxia encodes the alpha subunit of the hypoxia-inducible factor 1 transcription factor (HIF1α) [4]. HIF1 increases the expression of genes involved in diverse biological pathways [9, 10]. Many of the most prominent and well-characterised targets are involved in the regulation of oxygen supply and utilisation via angiogenesis and metabolic reprogramming [2, 9, 11]. Regulation of HIF activity has mainly been studied at the level of protein stability of the HIF1α subunit. HIF1α stability is regulated by several proline hydroxylases (PHDs), which modify proline residues in the oxygen-dependent degradation (ODD) domain of HIF1α [10, 12]. Hydroxylated HIF1α is recognised by the von-Hippel Lindau (VHL) E3-ubiquitin ligase, which promotes the ubuiquitination and subsequent degradation of HIF1α by the 26S proteasome[10, 12]. During hypoxia PHDs are inhibited resulting in rapid HIF1α accumulation, this allows HIF1α to dimerise with HIF1β to promote the expression of HIF target genes. [10, 12]. However, despite its much-increased half-life during hypoxia, studies have shown that HIF1α needs to be preferentially translated to sustain high HIF levels during prolonged hypoxic stress [4].

Translational control of individual mRNAs can be regulated by the non-coding regions at both the 5’ and 3’ ends of the transcript that can bind to regulatory proteins that control ribosome recruitment and translational initiation [5, 6, 8, 13]. Several studies using artificial reporter constructs have indicated that both the 5’ and 3’ UTRs of HIF1α can modulate translation rates through acting as internal ribosome entry sites (IRES), sites to recruit RNA binding proteins, or act as targets for miRNAs [13, 15]. However, the primary role for the endogenous HIF1α 5’UTR remains unclear, with conflicting studies using reporter constructs suggesting it represses translation through its complicated secondary structure [22], or activates translation by binding to specialised RNA-binding proteins to facilitate ribosome recruitment [15, 21]. As more recent work using genome editing approaches has indicated that reporter constructs investigating the role of UTRs do not always phenocopy the role of the endogenous UTRs in cells [23], we decided to modify the endogenous HIF1α 5’UTR in cells to investigate its function.

In this study we have investigated the consequences of uncoupling endogenous HIF1α from translational control in the cellular response to low oxygen in cell. Using a dual CRISPR/Cas9 strategy we determined the effects of deleting the 5’UTR of HIF1α in cultured cell lines. HIF1α 5’UTR deletion results in impaired hypoxia-induced HIF activity in cells. Furthermore, cells deleted for HIF1α 5’UTR demonstrate reduced solid tumour formation through impaired HIF1α translation rates. This study offers a genetic proof of principle that targeting HIF1α translation rates may be a strategy to suppress hypoxia-dependent gene expression in HIF-dependent human pathologies.

## MATERIALS AND METHODS

### Cell lines

PC-3 cells were grown in RPMI with 25mM HEPES, supplemented with 10% FBS and l-glutamine. U2OS and HeLa cells were grown in DMEM supplemented with 10% FBS, l-gluatamine, and penicillin streptomycin.

U2OS HRE-luciferase and HeLa HRE-luciferase were kind gifts from Professor Sonia Rocha and have been previously described [16]. PC-3 HRE luciferase cells were generated by co-transfection of 3xHRE-luciferase construct (Addgene 26371) with a puromycin-resistant construct. Stable clones were selected with 1.0 μg/ml and maintained in 0.5 μg/ml puromycin.

### CRISPR/Cas9 Genome Editing

CRISPR/Cas9 gRNAs were designed using zifit.partners.org to target regions within the HIF1α 5’UTR. Oligos 5’ CACCGATCACCCTCTTCGTCGCTT 3’ / 5’ AAACAAGCGACGAAGAGGGTGATC 3’ and 5’ CACCGCTTAGGCCGGAGCGAGCCT 3’/ 5’ AAACAGGCTCGCTCCGGCCTAAGC 3’ were annealed and cloned into the px458 cut with BsaI (Addgene #48138). Plasmids were transfected into cell lines using Genejuice transfection reagent (Merck) according to the manufacturer’s instructions. Clonal cell lines were isolated by limiting dilution with no antibiotic selection. Genomic DNA was isolated using DirectPCR lysis buffer (Viagen). Gene disruption was analysed by PCR using primers spanning the HIF1α 5’ UTR Sense 5’-cagtgctgcctcgtctga– 3’ and Antisense 5’ caatcccattaacgccgagg-3’. Sequencing reactions were performed using the primer 5’ caatcccattaacgccgagg 3’.

### Treatments

Cells were incubated at 1% O_2_ in an *in vivo* 300 hypoxia workstation (Ruskin, UK). Cells were lysed for protein extracts and RNA extraction in the chamber to avoid re-oxygenation.

### Cell lysis and immunoblotting

Cell lysis and immunoblotting was performed as described [17, 18]. Briefly, cells were lysed in 8 M urea lysis buffer or RIPA buffer and immunoblotted as described. Antibodies used were HIF1α (Clone 241809, R&D systems), HIF2α (#7096, Cell Signaling Technologies), YB-1 (A303–231A, Bethyl), YB-1 (#4202, Cell Signaling Technologies), BNIP3L (#12396, Cell Signaling Technologies), NDRG1 (#5196, Cell Signaling Technologies), β-actin (AC-74, Sigma).

### Luciferase Assays

Lysates for luciferase assay were prepared in 1× passive lysis buffer (Promega), 100 μl per well of a 24-well plate. 10 μl of lysate was incubated with 50 μl luciferase reagent (Promega) and measured for 10 s using Lumat LB9507 (EG&G Berthold). Graphs represent raw Relative Luminescence Unit (RLU) readings from three independent experiments.

### Polysome profiling

Cells were grown to 80% confluency and incubated in 100 mg/ml cycloheximide for 3min and resuspended in hypotonic polysome extraction buffer (5 mM Tris [pH 7.5], 2.5 mM MgCl_2_, 1.5 mM KCl, 1% Triton X-100, 100 mg/ml cycloheximide, 100 U/ml RNasin). Cells were lysed through the addition of Triton X-100 (0.5%) and sodium deoxycholate (0.5%) to solubilise the cytosolic and endoplasmic reticulum-associated ribosomes. Extracts were normalised by OD 260 nm and layered onto 10 ml of 10– 50% sucrose steps and centrifuged at 222 228 × g (36 000 rpm) for 2h at 4°C using SW41Ti rotor. The sucrose steps were manually fractionated into twelve 0.75 ml fractions. Absorbance at OD_254 nm_ and visualisation by RNA agarose electrophoresis following Trizol (Invitrogen) purification was used to determine the monosomal and polysomal fractions. Polysomal fractions were pooled for subsequent qRT-PCR analysis.

### RNA immunoprecipitation

To probe for direct interactions between YB-1 protein and HIF1α transcripts, PC-3 were resuspended in hypotonic polysome extraction buffer and lysed through the addition of Triton X-100 (0.5%) and sodium deoxycholate (0.5%) as described in [17]. Clarified lysates were incubated with 1 ug of YB-1 antibody and antibody/protein/RNA complexes were isolated by using 20 μl packed volume protein A sepharose beads (Invitrogen). Beads were washed with polysome extraction buffer and RNA isolated using the PeqGold RNA isolation kit. cDNA was prepared using Qiagen Quantanova cDNA synthesis kit.

### Quantitative reverse transcription-PCR

Quantitative PCR data was generated on a Rotor-Gene Q (Qiagen) (or stratagene) using the following experimental settings: hold 50°C for 3 min; hold 95°C 10 min; cycling (95°C for 30 s; 58°C for 30 s; 72°C for 30 s with fluorescence measurement for 45 cycles). All values were normalised to 18S rRNA or RPL13A levels using the Pfaffl method as indicated. Primers sequences: RPL13A sense 5′-CCT GGA GGA GAA GAG GAA AGA GA-3′, antisense 5′-TTG AGG ACC TCT GTG TAT TTG TCA A-3′; BNIP3L sense 5′-GTC GCC TGT CCA CTT AGC C-3′, antisense 5′-GCT GTT TGC CCG TTC TTA TTA CA-3′; NDRG1 sense 5′-CTG GCA TCA ACG CTG TCT TC-3′, antisense 5′-GCC TAT GAG GTG CAG GGT C-3’; HIF1α coding For-5′-CATAAAGTCTGCAACATGGAAGGT-3′, HIF1α Rev 5′-ATTTGATGGGTGAGGAATGGGTT-3′; 18S rRNA For 5′-GTAACCCGTTGAACCCCATT-3′, 18S rRNA Rev 5′-CCATCCAATCGGTAGTAGCG-3′;

### Quantitation of HIF1α transcript levels

Quantitation of HIF1α transcript levels. The primers designed to recognise individual HIF1α mRNA variants are as follows: HIF1α mRNA NM_001530.4 (P1) sense 5’-TTTCCTTCTCTTCTCCGCGT-3’, antisense 5’-CCCTCCATGGTGAATCGGT-3’; HIF1α mRNA NM_181054.2 (P1&2) sense 5’-GGGACCGATTCACCATGGA-3’, antisense 5’-CTTTACTTCGCCGAGATCTGG-3’; HIF1α mRNA NM_001243084.1 (P3) sense 5’-GGGACCGATTCACCATGGA-3’, antisense 5’-CTTTACTTCGCCGAGATCTGG-3’, HIF1α all isoforms For-5′-CATAAAGTCTGCAACATGGAAGGT-3′, HIF1α Rev 5′-ATTTGATGGGTGAGGAATGGGTT-3′. Inverse log Ct was calculated using the efficiency values for each primer set and normalised to the total HIF1α levels using the value from the HIF1α total qRT-PCR primer set. Experiments were performed twice with 3 biological replicates for each individual cancer cell line for each qRT-PCR.

### Cell Proliferation Assays

For each cell line 1 ×10^5^ cells were seeded per well in 6-well plates. Cells were trypsinised and counted using a haemocytometer after 1, 2, 3 and 4 days cell culture. For hypoxia treatments cells were allowed to attach to plates for 24 hours before being transferred to the hypoxia workstation. Mean and standard errors are displayed and area under the curve calculated as a measure of cell proliferation.

### Clonognenic Assays

Clonogenic assay. CRISPR/Cas9-modified PC-3 cells were cultured in normoxia or hypoxia for 24h. Cells were trypsinised and plated in 6 well plates at a concentration of 200 cells per well. Plates were incubated at 21% O_2_ to allow colony formation for 14 days. The colonies were fixed with Carnoy’s fixative (3:1 methanol:acetic acid) and stained with 0.4% (w/v) crystal violet. Finally, the plates were inspected by microscopy and the number of the colonies were counted. Each assay was made in triplicate and only colonies containing at least 50 cells were counted. Colony formation was normalised to plating efficiency calculated for each cell line.

### Ethics statement

All mouse experiments were approved by Newcastle University’s Animal Welfare and Ethical Review Board. All procedures were carried out under project and personal licenses approved by the Secretary of State for the Home Office, under the United Kingdom’s 1986 Animal (Scientific Procedures). Animals were housed in the Comparative Biology Centre, Newcastle University animal unit.

### Sub-cutaneous xenograft studies

NOD-SCID mice were implanted sub-cutaneously with 1 × 10^7^ PC3 prostate cancer cells that were either HIF1-alpha WT or genetically modified to have part of the 5’-untranscribed region (5’-UTR) of HIF1-alpha missing. Cells were implanted into the hind flank and the tumour growth and burden was monitored at least three times weekly using calipers. Mice did not undergo any anaesthesia at any point during this study. The endpoint for this study was pre-defined at a PC3 tumour volume of 400mm^2^ (20 mm in any direction by caliper measurement) and mice were humanely killed by the Schedule 1 method of cervical dislocation at this point. It is important to note that this experimental work was performed following the blinding of the genotype of the cells prior to implantation.

### Immunohistochemistry

Formalin-fixed tumour tissues were paraffin-embedded and serial sections were cut by the Molecular Pathology Node, Cellular Pathology, Royal Victoria Infirmary, Newcastle-Upon-Tyne.

Formalin-fixed paraffin-embedded tumour sections were dewaxed and hydrated. Endogenous peroxidase activity was blocked with hydrogen peroxide and antigen retrieval was achieved using 1 mM EDTA. Tissue was blocked using an Avidin/Biotin Blocking Kit (SP-2001, Vector Laboratories, Peterborough, UK) followed by 20% swine serum in PBS and then incubated with primary antibodies overnight at 4 °C (γH2AX ((#9718, Cell Signaling Technologies). The following day, slides were washed and incubated with biotinylated swine anti-rabbit (E0353, Dako, UK) followed by Vectastain Elite ABC Reagent (PK7100, Vector Labs). Antigens were visualised using DAB peroxidase substrate kit (SK4001, Vector Labs) and counterstained with Mayer’s haematoxylin. Immuno-stained cells were imaged using a DFC310 FX microscope (Leica Microsystems) and the images blinded (coded) prior to analysis. At least 3 images per tissue at × 100 magnification (10x lens and 10x eye piece) were analysed using brown/blue pixel intensity using Adobe Photoshop.

## RESULTS

### High efficiency modification of the HIF1α 5’UTR using a dual CRISPR/Cas9-based approach

The activity of the HIF1 transcription factor is controlled primarily by the availability of the HIF1α subunit, which is rapidly stabilised during hypoxia via the PHD/VHL axis [2, 11]. However, even in hypoxic conditions HIF1α remains relatively unstable, with a half-life of approx. 60-90min at 1% O_2_ in PC-3 prostate cancer cells, indicating that active translation of the HIF1α transcript is necessary to maintain hypoxia-induced HIF activity (Supplemental Figure 1A and B). This highlights that control of HIF1α protein translation rates is therefore an important additional mechanism to control HIF activity in hypoxic cells [4, 8, 17, 19-21]. To investigate the role of the endogenous HIF1α 5’UTR the HIF1α transcript variants were quantified in prostate cancer (PC-3), osteosarcoma (U2OS) and cervical cancer (HeLa). qRT-PCR analysis revealed that the two most predominant HIF1α transcripts in PC-3, Hela, and U2OS cell lines are encoded from the same exon, but from alternative transcription start sites encoding HIF1α mRNAs containing a 292bp 5’UTR and a 404bp 5’UTR (Supplemental Figure 2A). An annotated splice variant of HIF1α mRNA encoding a 229bp 5’UTR was virtually absent from any of the cell lines examined (Supplemental Figure 2B). To assess the role of the endogenous HIF1α 5’UTR, a CRISPR/Cas9-based genome editing strategy using dual gRNAs was used to delete a segment of the 5’UTR of both 292bp 5’UTR and a 404bp 5’UTR without deleting either transcription start site (Figure 1A). Dual CRISPR gRNAs and Cas9 transiently transfected into PC-3, HeLa, and U2OS cells resulted in a detectable and predictable deletion of the HIF1α 5’UTR as measured by semi-quantitative PCR (Figure 1B-D). qRT-PCR revealed CRISPR/Cas9-dependent modification reduced the number of HIF1α mRNAs containing the 5’UTR (Figure 1E-G), without significantly altering total HIF1α transcripts (Figure H-J).

**Figure 1.**
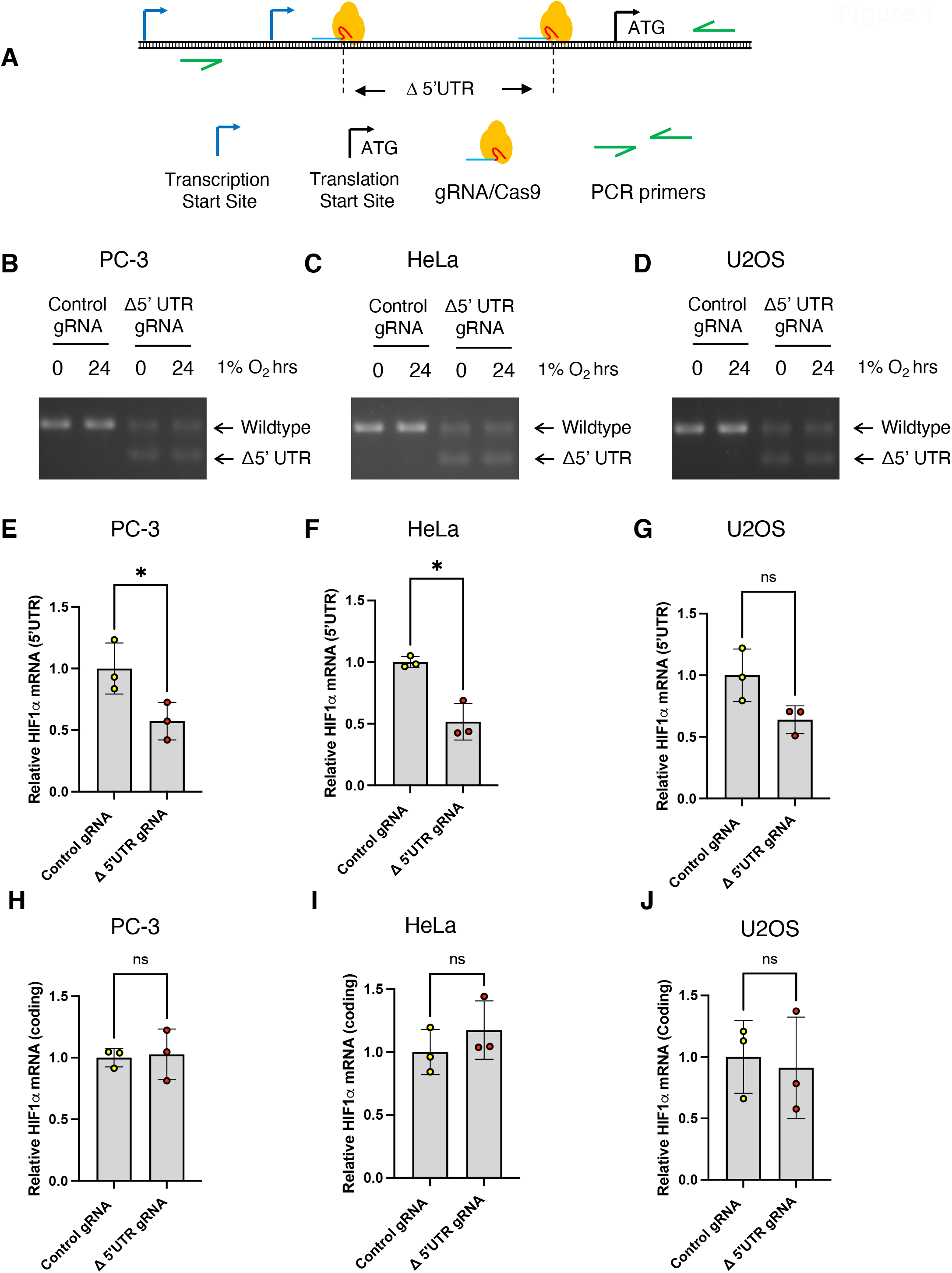
Efficient and predictable deletion of the HIF1α 5’UTR using a dual CRISPR gRNA strategy. **(A)** Schematic of CRISPR/Cas9 experimental design and PCR analysis of genomic DNA encoding the HIF1Α 5’UTR. Transcription start sites for HIF1α mRNA v1 and v2 are indicated in addition to the common translation start side. (**B-D)** PCR analysis of genomic DNA isolated from (B) PC-3, (C) HeLa or (D) U2OS, cells transfected all transfected with Cas9, and a control gRNA or the HIF1α 5’UTR gRNAs. The arrows indicate the size of the PCR product from the unmodified cells (Wildtype) and the predicted 5’UTR deletion (Δ5’ UTR). (**E-G)** Quantitative RT–PCR analysis of 5’UTR HIF1α mRNA prepared from (E) PC-3, (F) HeLa or (G) U2OS cells transfected with Cas9 and a control gRNA, or a pair of gRNAs designed to delete the HIF1α 5’UTR. 5’UTR HIF1α mRNA values normalized to HIF1α coding region. **(H-J)** Quantitative RT– PCR analysis of HIF1α mRNA prepared from (H) PC-3, (I) HeLa or (J) U2OS cells transfected with Cas9 and a control gRNA, or a pair of gRNAs designed to delete the HIF1α 5’UTR. HIF1α coding region values are normalized to RPL13A mRNA and fold change calculated from control samples. Statistical analysis performed using a Students T-test.

**Figure 2.**
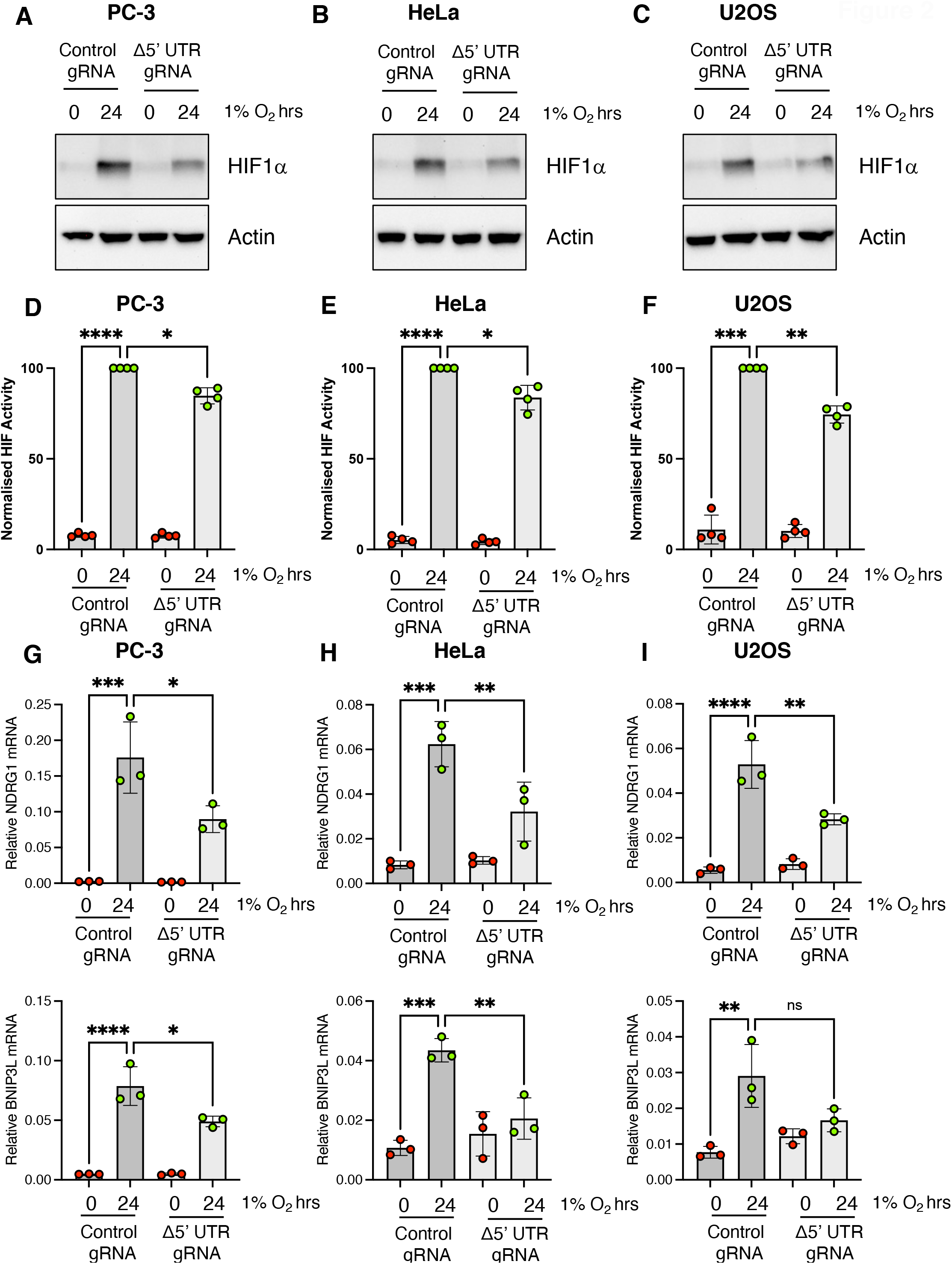
Deletion of the HIF1α 5’UTR by transient transfection impairs HIF1α accumulation and HIF1 activity in hypoxic tumour cells. **(A)** PC-3, **(B)** HeLa and **(C)** U2OS cells transfected with Cas9, and a control gRNA or HIF1α 5’UTR gRNAs. Cells were exposed to hypoxia as indicated and WCLs subjected to immunoblot analysis to assess expression levels of the indicated proteins. **(D)** U2OS, **(E)** HeLa and **(F)** PC-3 cells stably expressing luciferase driven from the hypoxia responsive element (HRE) were transfected with Cas9 and the indicated gRNAs and exposed to 1% O_2_ for 24h. Results presented represent the mean plus S.D. of four experiments. Quantitative RT–PCR analysis of NDRG1 and BNIP3L mRNA prepared from **(G)** PC-3, **(H)** HeLa and **(I)** U2OS cells transfected with Cas9 and the indicated gRNAs and exposed to 1% O_2_ for 24h. All values are normalised to RPL13A mRNA. Statistical analysis performed using a one-way ANOVA with Dunnett’s multiple comparisons test.

### HIF1α 5’UTR deletion reduces HIF1α levels and activity in cells

As discussed above, the 5’UTR of HIF1α mRNA has been proposed to have conflicting roles in controlling HIF1α translation rates in cells [15, 17, 24]. As a significant number of HIF1α mRNAs were modified in cells transiently transfected with dual CRISPR gRNAs and Cas9, we assessed the effect of endogenous HIF1α mRNA 5’UTR deletion on hypoxia-induced HIF1α protein levels in the mixed pooled population. Immunoblot analysis revealed robust hypoxia-induced HIF1α accumulation in PC-3, U2OS, and HeLa cells (Figure 2A-C). However, HIF1α accumulation was attenuated in CRISPR/Cas9-modified populations of PC-3, U2OS, and HeLa cells containing cells missing the HIF1α mRNA 5’UTR, consistent with the 5’UTR playing a positive role in maintaining high HIF1α levels in response to hypoxic stress (Figure 2A-C). HIF activity was then assessed by transiently expressing CRISPR/Cas9 gRNAs designed to remove the 5’UTR of HIF1α mRNA into PC-3, HeLa, and U2OS cells containing an integrated luciferase reporter construct possessing three copies of the hypoxia-responsive element (HRE) consensus-binding site (Figure 2D-F). A robust activation of luciferase activity was observed in cells exposed to 1% O_2_, which was weakly but significantly reduced in cells transfected with 5’ UTR-targeting CRISPR/Cas9 constructs (Figure 2D-F). The effect of HIF1α 5’UTR deletion on endogenous HIF1 target gene expression was then assessed by quantitative RT-PCR of N-myc downregulated gene 1 (NDRG1) and BCL2 Interacting Protein 3 Like (BNIP3L) mRNAs, both of which are endogenous HIF1 target genes [25, 26]. Both NDRG1 and BNIP3L mRNA levels were induced in hypoxic PC-3, HeLa, and U2OS cells, as expected (Figure 2G-I). Transient expression of CRISPR/Cas9 gRNAs designed to remove the 5’UTR of HIF1α mRNA reduced hypoxia-induced expression of both BNIP3L and NDRG1 mRNAs, consistent with a reduction in HIF1 activity in modified cells (Figure 2G-I). Together these data indicate the endogenous 5’UTR of HIF1α mRNA plays a general role in maintaining high HIF1α levels during hypoxia in multiple cell types derived from different human tissues.

### The 5’UTR of HIF1α mRNA is required to maintain HIF1α mRNA translation during hypoxic stress

To allow detailed analysis of the contribution of the HIF1α mRNA 5’UTR on the HIF-dependent hypoxic response we isolated monoclonal PC-3 prostate cancer cell lines homozygous for the HIF1α 5’UTR deletion (Supplemental Figure 3A). Deletions were confirmed by PCR and defined by Sanger sequencing of the PCR products (Supplemental Figure 3A-B). Intriguingly, deletions from independent clonal cell lines had the same 195bp deletion, corresponding to a single major deletion product observed in the transient transfections with the dual gRNAs, indicating a consistent mechanism of DNA repair (Figure 1B-D, Supplemental Figure 3B).

**Figure 3.**
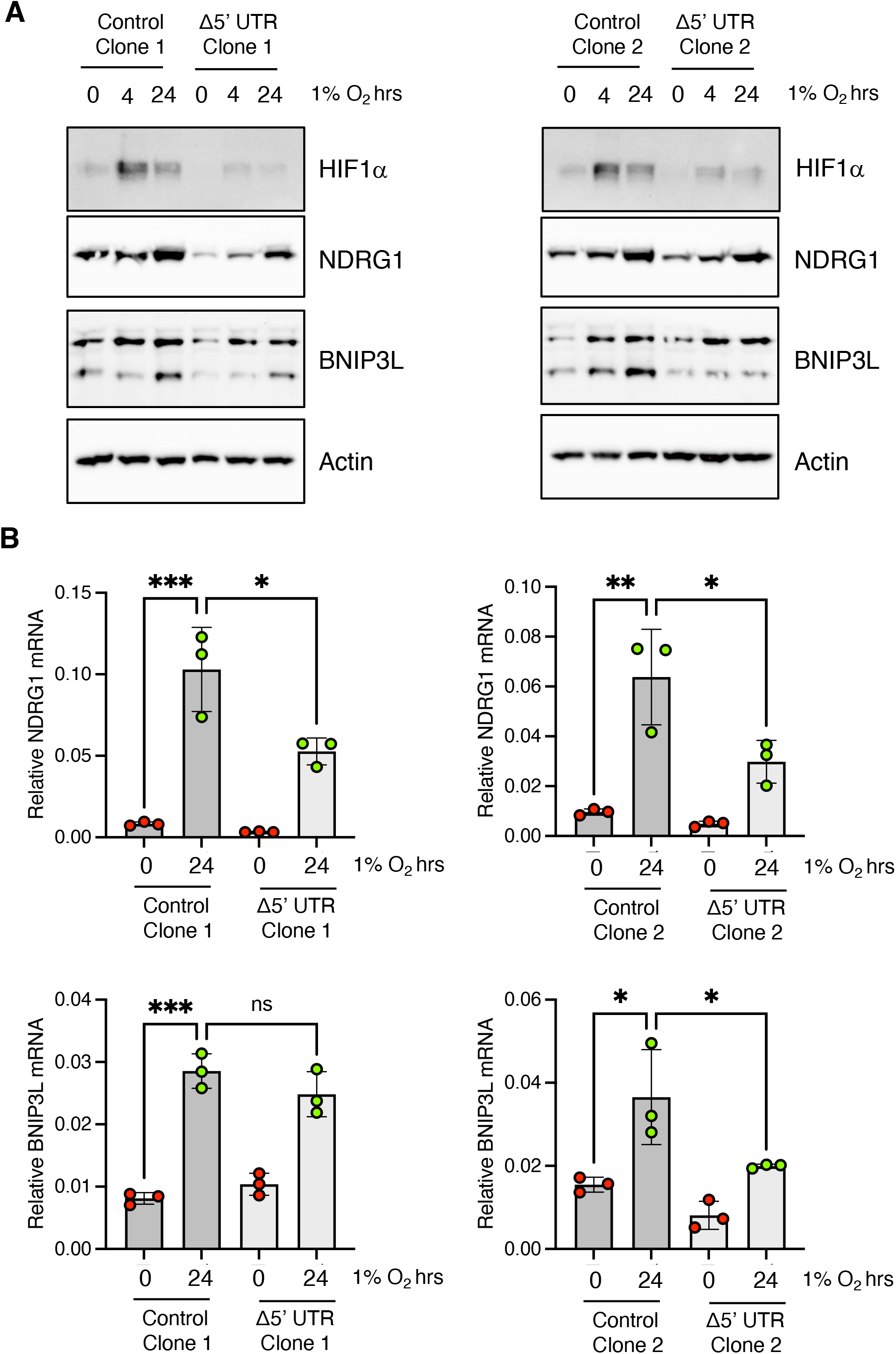
HIF1α 5’UTR deletion in clonal cell lines have impaired HIF1-depedent gene expression. **(A)** WCLs were prepared from PC-3 derived clonal cell lines, wildtype or containing the HIF1α 5’UTR deletion, exposed to 1% O_2_ as indicated. WCLs were resolved by SDS PAGE and analysed by immunoblot using the indicated antibodies. **(B)** Quantitative RT– PCR analysis of NDRG1 and BNIP3L mRNA prepared from PC-3 clonal cell lines exposed to 1% O_2_ for 24h. All values are normalised to RPL13A mRNA.

To examine if the reduced HIF1α phenotype was maintained after clonal selection, hypoxia-induced HIF1α accumulation during hypoxia was measured by immunoblot (Figure 3A). As observed when cells were transiently transfected, 2 independent HIF1α 5’UTR deletion clones had an impaired level of HIF1α accumulation in response to hypoxic stress (Figure 3A). Similarly, levels of hypoxia-induced NDRG1 and BNIP3L were reduced at both the protein and mRNA level indicating that the HIF1α 5’UTR is necessary for the full HIF1-dependent hypoxic response (Figure 3A and 3B). These data show that during clonal selection there was no compensatory adaptation to reverse defective HIF1α accumulation following hypoxic stress (Figure 3A and 3B).

Polysome profiling was then performed to examine the effect of HIF1α mRNA 5’UTR deletion on protein translation rates in response to low oxygen. A significant decrease in global translational efficiency was observed in wildtype PC-3 cells exposed to hypoxic stress, as indicated by the large increase in mRNAs associated with monosomal fractions, and a concomitant decrease in the number of mRNAs associated with polysomes (Figure 4A). A similar block in global protein synthesis rates was observed in cells missing the HIF1α 5’UTR indicating the 5’UTR of HIF1α mRNA has no overt effect on global protein translation rates (Figure 4A). To investigate the levels of individual mRNAs associated with actively translating ribosomes, cDNAs were prepared from fractions containing polysomes (Fractions 8– 11) and compared to relative quantities of mRNAs in the input samples. Quantitative RT-PCR analysis of polysome-associated HIF1α mRNA revealed that HIF1α translation rates increased in hypoxic wildtype PC-3 cells, compared to normoxic cells, confirming that HIF1α mRNA is preferentially translated during hypoxic stress (Figure 4B). In contrast, translation rates of HIF1α in Δ5’UTR PC-3 cells were markedly impaired during hypoxia, demonstrating that the 5’UTR plays an important role in directing preferential HIF1α mRNA translation during hypoxia stress (Figure 4B). These data suggest that genetic deletion of the 5’UTR uncouples HIF1α mRNA from translational control.

**Figure 4.**
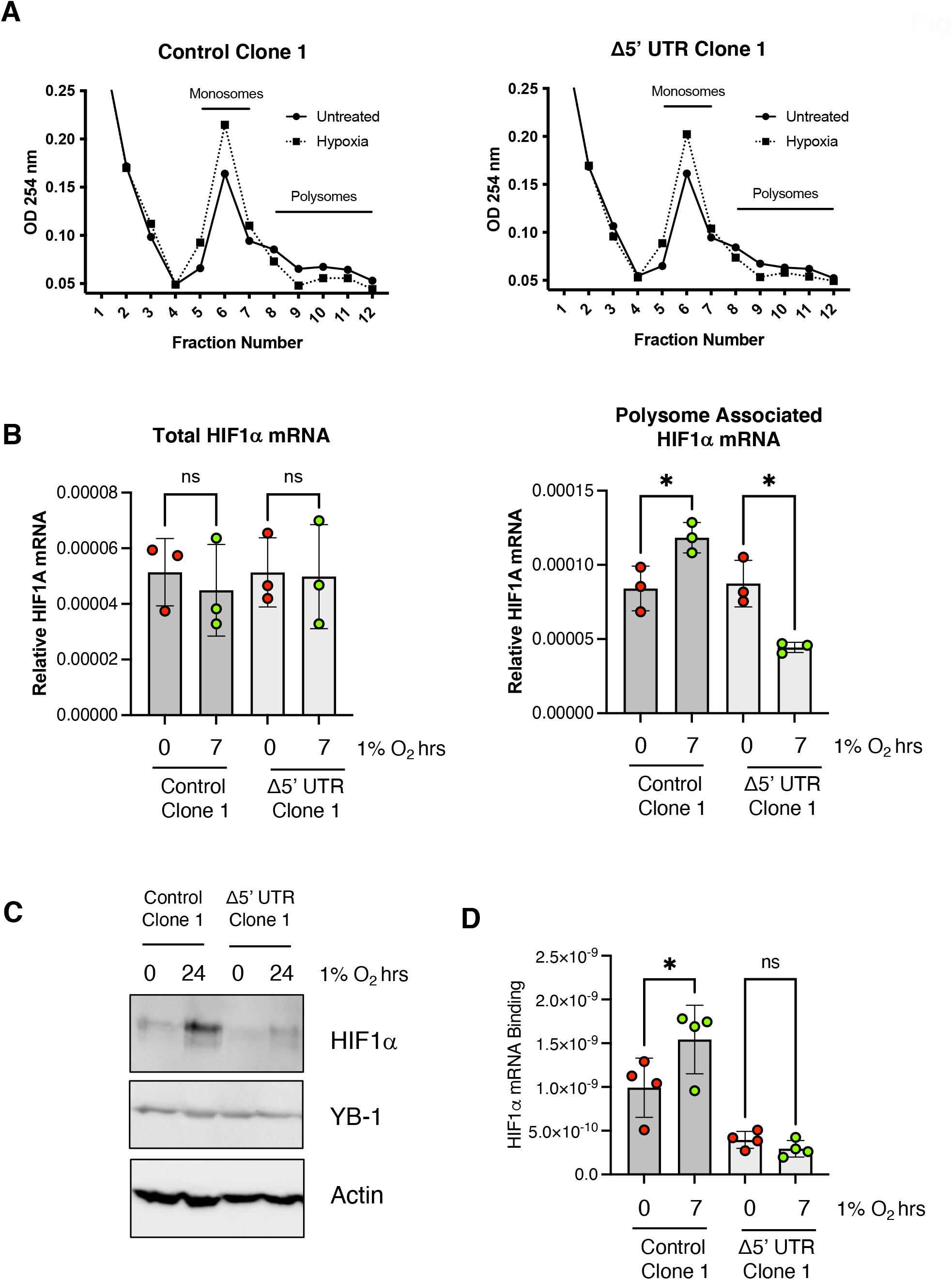
HIF1α 5’UTR deletion suppresses HIF1α translation by preventing YB-1 binding to HIF1α mRNA. **(A)** Representative polysome profiles of WT and D5’UTR PC-3 cells exposed to 1% O_2_ for 7 h. Fractions containing monosomes and polysomes are indicated on the graph. **(B)** Quantitative RT-PCR analysis of HIF1α mRNA prepared from input mRNA and polysomal fractions (8–11) prepared from WT and D5’UTR PC-3 cells exposed to 1% O_2_ for 7 h as indicated. All values are normalised to 18S rRNA. (C) WCLs were prepared from PC-3 derived clonal cell lines, wildtype or containing the HIF1α 5’UTR deletion, exposed to 1% O_2_ as indicated. WCLs were resolved by SDS PAGE and analysed by immunoblot using the indicated antibodies. **(D)** WCLs were prepared from PC-3 derived clonal cell lines, wildtype or containing the HIF1α 5’UTR deletion, exposed to 1% O_2_ as indicated. WCLs were resolved by SDS PAGE and analysed by immunoblot using the indicated antibodies. **(E)** HIF1α mRNA bound to YB-1 in WT and D5’UTR PC-3 cell lines in the presence of 1% O_2_ as indicated. Significance was calculated relative to untreated control using a one-way ANOVA using the Dunnett multiple comparison test.

### HIF1α mRNA/YB-1 interaction is disrupted in Δ5’UTR cells

The 5’UTR of several stress-inducible factors can bind to RNA-binding proteins which alter translation rates by modulating ribosome recruitment [4, 5, 13, 14, 27]. Several regulatory factors have been identified that enhance or reduce HIF1α translation by directly binding to the HIF1α mRNA 5’UTR[4, 15]. We and others have previously described a central role for the Y-box binding protein 1 (YB-1) in binding to the HIF1α 5’UTR and promoting HIF1α translation rates [17, 21, 28]. Immunoblot analysis of YB-1 levels from PC-3 WT and Δ5’UTR cells either cultured in normal oxygen conditions or hypoxia revealed no significant changes in total YB-1 levels between cell lines, or in response to hypoxic stress (Figure 4C). As total levels of YB-1 are unchanged, we investigated if the YB-1/HIF1α mRNA interaction is disrupted by deletion of the HIF1α 5’UTR. cDNA prepared from YB-1 precipitates and analysed by qRT-PCR revealed an increased association between YB-1 and HIF1α mRNA in hypoxia is supported by previous observations [17] and highlights its role in directing hypoxia-induced HIF1α translation rates (Figure 4D). Cells expressing HIF1α mRNA lacking the 5’UTR lost the HIF1α mRNA/YB-1 interaction, which was no longer induced by hypoxia (Figure 4D). The reduction of the HIF1α mRNA/YB-1 interaction is consistent with the observed reduction in hypoxia-induced HIF1α mRNA translation rates observed in the Δ5’UTR cell lines and indicates that key regulatory protein interaction sites are removed from HIF1α mRNA missing the 5’UTR (Figure 4B).

### Deletion of the HIF1α 5’UTR sensitises cells to hypoxic stress

Genetic deletion of the endogenous HIF1α 5’UTR allows analysis of the cellular consequences of disrupting translational control of HIF1α *in vivo*. In normal oxygen conditions, deletion of the HIF1α UTR did not alter cellular proliferation rates consistent with cells remaining well oxygenated (Figure 5A and 5B). However, when proliferation rates were measured in cells grown at 1% O_2_, Δ5’UTR cells had a markedly reduced proliferation rate, as compared to wildtype cells (Figure 5C and 5D). This suggests that the reduced HIF1 activity in Δ5’UTR cells has a negative effect on proliferation rates, and/or cellular viability in cells grown in hypoxic conditions (Figure 5C and 5D). Subsequently, we investigated whether deletion of the 5’UTR of HIF1α could impair PC-3 cell colony-forming ability after hypoxic stress. Wildtype or Δ5’UTR PC-3 cells were cultured in normoxic conditions or 24h in 1% O_2_ and then seeded at low density to examine colony formation ability. Removal of the HIF1α 5’UTR had no significant effect on PC-3 cells cultured in normoxia to form colonies, however cells missing the 5’UTR of HIF1α mRNA had a significantly reduced ability to form colonies following hypoxic stress (Figure 5E). These data suggest that removal of the 5’UTR of HIF1α mRNA from cells reduces viability in response to hypoxic stress.

**Figure 5.**
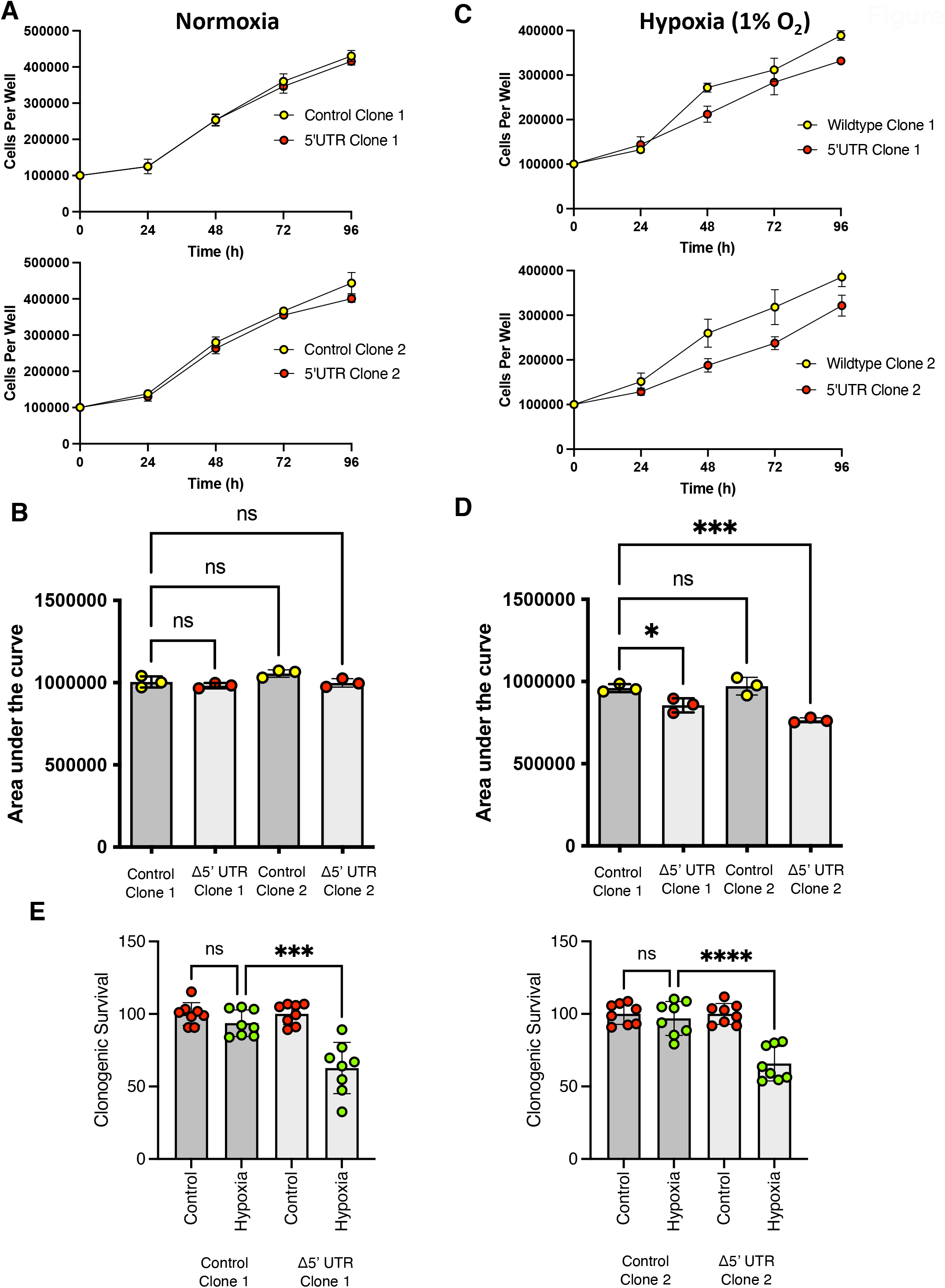
HIF1α 5’UTR deletion sensitises cells to hypoxic stress. **(A)** WT or 5’UTR clonal cell lines were plated out on 6 well plates at a seeding density of 100,000 cells per well. Cells were trypsinised and counted after days 1, 2, 3 and 4. **(B)** Area under the curve was calculated from data in (A) to determine differences in growth rates between clonal cell lines **(C)** WT or 5’UTR clonal cell lines were plated out on 6 well plates at a seeding density of 100,000 cells per well. Cells were left to attach for 6 hours and transferred to 1% O_2_. Cells were trypsinised and counted after days 1, 2, 3 and 4. **(D)** Area under the curve was calculated from data in (C) to determine differences in growth rates between clonal cell lines **(E)** Clonogeneic survival of WT or D’ 5’UTR deleted cell lines pre-exposed to hypoxic stress before being plated out at 200 and 400 cells per 6 well plates. Colonies of over 50 cells were counted. Colony counts were normalised to the plating efficiency of Clone 1 WT PC-3 cells. Significance was calculated relative to untreated control using a one-way ANOVA using the Dunnett multiple comparison test.

### 5’UTR deletion reduced tumour formation in tumour models

In solid tumours activation of hypoxia-responsive signalling pathways contributes to the malignant phenotype by promoting the expression of pro-angiogenic and pro-survival gene products [1, 2, 29, 30]. As such, hypoxia, and hypoxia-induced signalling pathways are often hijacked by the tumour cells to survive and thrive in the hypoxic tumour micro-environment [2,11]. To test the role of the endogenous HIF1α 5’UTR on the formation of solid tumours *in vivo* we implanted WT and Δ5’UTR PC-3 cells into the flank of nude mice. Tumour volume was measured by callipers over a 3-week period. Both wildtype and Δ5’UTR PC-3 cells initially form tumours at equivalent rates (Figure 6A) and all mice maintained consistent, healthy bodyweights (Supplementary Figure 4)). However, approximately 12-14 days post implantation, Δ5’UTR PC-3 derived tumours started to regress, while WT tumours keep growing (Figure 6A). Upon study termination, both tumour volume and weight (Figure 6B-D) confirmed that WT tumours were larger than those containing Δ5’UTR PC-3 cells, with one mouse appearing to have undergone a complete tumour regression. Interestingly, we also found that DNA damage (as measured by γH2AX staining) was increased in the WT PC3 tumours, although the difference was not statistically significant (Figure 6E). This may suggest that these tumours are more genomically unstable and have a higher rate of cell turnover than the cells within the Δ5’UTR PC-3, which would be consistent with continued growth of the WT tumours but not the Δ5’UTR PC-3 counterparts.

**Figure 6.**
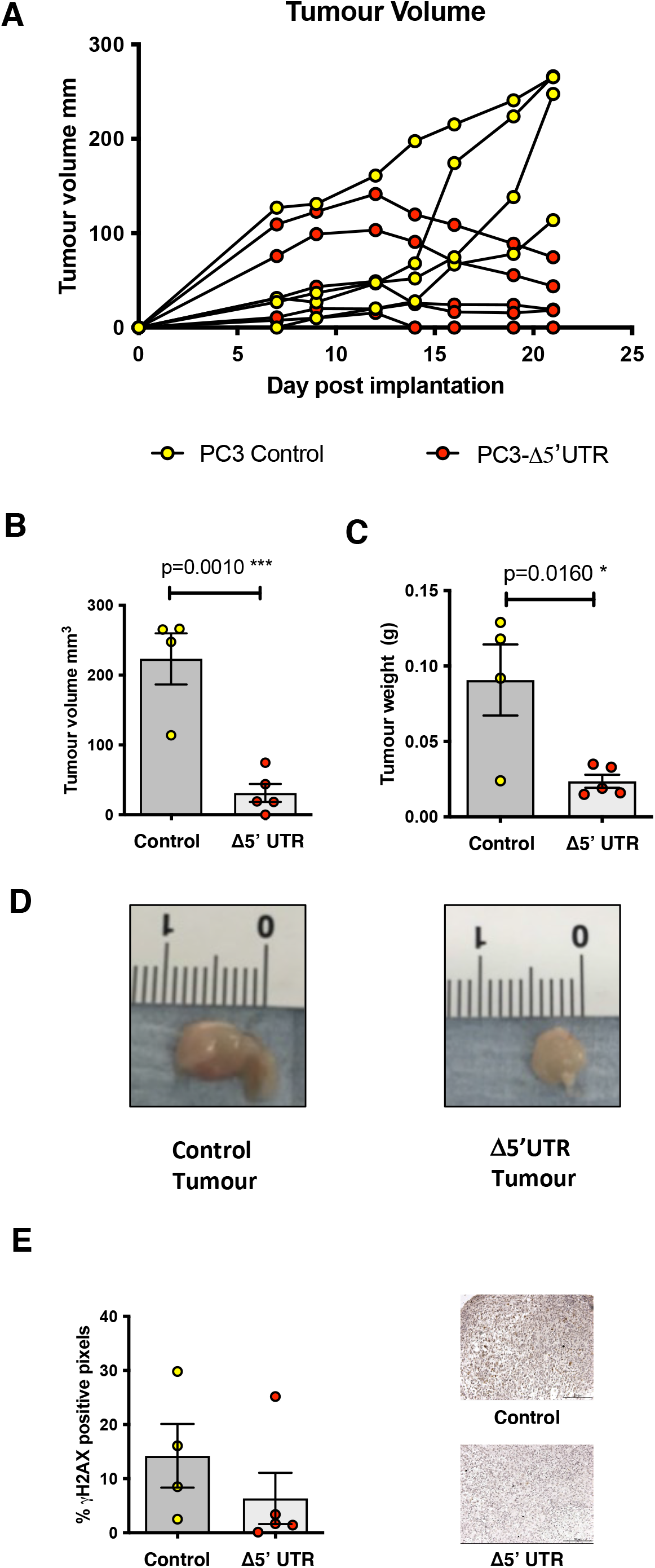
Defective HIF1α protein translation limits tumour growth in vivo. **A)** WT PC-3 cells or 5’UTR PC-3 cells xenografts were implanted into nude mice. Tumour volume was measured by calipers and plotted. (B) Tumour Volume (C) Tumour Weight (D) Representative Images.

Collectively our data reveal that the 5’UTR of the HIF1α mRNA is necessary for full activation of the HIF-dependent hypoxic response and solid tumour formation in vivo. Our data suggest that suppression of HIF1α translation may provide a novel strategy for targeting aberrant HIF activity in human disease.

## Discussion

Disruption to oxygen homeostasis and aberrant HIF activity is observed in multiple human pathologies such as heart disease, cancer, cardiovascular disease and chronic obstructive pulmonary disease[1, 2, 11, 29]. Most studies examining how HIF is deregulated have focused on protein degradation, however the contribution of HIF1α translation rates, and the underlying mechanistic control is rarely considered [4-6, 31]. The results from the present study, for the first time, directly measure the contribution of targeted control of mRNA translation in the regulation of the HIF-dependent hypoxic response in both 2D cell culture and the formation of solid tumours. Our work clearly demonstrates the importance of HIF1α translation control for its function.

Translational regulation of mRNA is an understudied, yet critically important step in the control of gene expression [6, 32]. A number of stresses including starvation, heat/ cold shock, DNA damage, and hypoxia, lead to changes in gene expression patterns caused by a general shutdown and reprogramming of protein synthesis [6]. However, cells have mechanisms to ensure mRNAs from stress-responsive genes are translated to restore homeostasis [5-8, 13, 31]. In this study we interrogated the role for the 5’UTR of the HIF1α mRNA in regulating HIF1α mRNA translation rates during hypoxia. We found that CRISPR/Cas9-dependent deletion of the 5’UTR of the HIF1α mRNA removes HIF1α mRNA from hypoxia-dependent translational control, resulting in attenuated HIF-dependent gene expression. HIF1α mRNA lacking this 5’UTR is no longer efficiently translated in hypoxic cells, leading to impaired HIF activity and reduced cell survival during hypoxic stress. Cell unable to preferentially translate HIF1α in response to hypoxia are unable to form large solid tumours in murine xenografts.

The translation machinery can respond to an array of cis-acting elements, located on the RNA transcript, which dictate the fate of mRNAs [13, 14]. These include not only the 5’ UTR, but also the 3’UTR and a variety of post-transcriptional modifications that can ultimately contribute to translation rates [13]. Our work has shown that the 5’UTR of HIF1α mRNA is necessary for the HIF-dependent hypoxic response by regulating HIF1α mRNA association with polysomes and regulatory proteins such as YB-1. In addition other work has shown the 3’ UTR of HIF1α also plays a positive role in regulating HIF1α mRNA translation rates by recruiting several translation initiations factors, including eiF3 and DHX29, which unwind the 5’UTR during translation initiation [22]. This work indicates that the 5’ and 3’ HIF1α UTRs act in concert to control HIF1α protein translation, indicating that examining the roles of UTRs in the context of an artificial reporter construct may miss the full picture of regulation [23].

Proof of principle studies using siRNA and genetic deletion of HIF1α showed that reducing HIF1 activity in solid tumours inhibits primary tumour growth [33-35]. As such chemical manipulation of the HIF pathway by small molecule modulators is widely perceived as a promising strategy to target diseases associated with HIF dysfunction [29, 30, 36, 37]. Compounds that attenuate HIF1α translation rates have been shown to have anti-angiogenetic and anti-tumour activity, but due to their mechanism of action are likely to have many unwanted on-target and off-target effects [38-41]. Our work using a tractable genetic model gives insights into both the mechanisms behind HIF1α translational control, and the consequences of disrupting it. Targeting translation factors that control hypoxia dependent gene expression, such as YB-1, is an efficient strategy to suppress HIF activity during solid tumour growth and development. Indeed, a recent study demonstrated that YB-1 acetylation could limit tumour metastasis, in part by blocking translational activation of HIF1α and other cytoprotective factors [8]. The authors of this study demonstrated that HDAC inhibition increased levels of YB-1 acetylation, and decreased tumour size and metastatic potential in murine xenografts [8].

Collectively our data reveal a critical role for the endogenous 5’UTR of the HIF1α mRNA in the regulation of HIF1α translation rates during hypoxic stress. Strategies to suppress HIF1α translation rates to specifically inhibit HIF1α levels may provide a novel strategy for targeting aberrant HIF activity in human disease, while still maintaining essential normal cell function.

## Acknowledgements

Funding for this work was provided by North West Cancer Research Development Grant [RDG2021.12 to NSK], Cancer Research UK [C1443/A22095 to JEH/ NDP];.BBSRC DTP Studentship [BB/T008695/1 to GBE] and Wellcome Trust [206293/Z/17/Z to FC and SR]

## Author contributions

JEH and NSK designed the research. JEH, OM, GE, FC, HM, GR, MD and NSK performed the research. JEH, OM, GE, FC, HM, GR, MD, SR, NDP and NSK analysed the data. NSK wrote the manuscript. JEH, SR, NDP and NSK edited the completed manuscript.

