## Supplemental Figures for "Disruption of HIF1Α translational control attenuates the HIF-dependent hypoxic response and solid tumour formation *in vivo*"

### Supplemental Figure 1

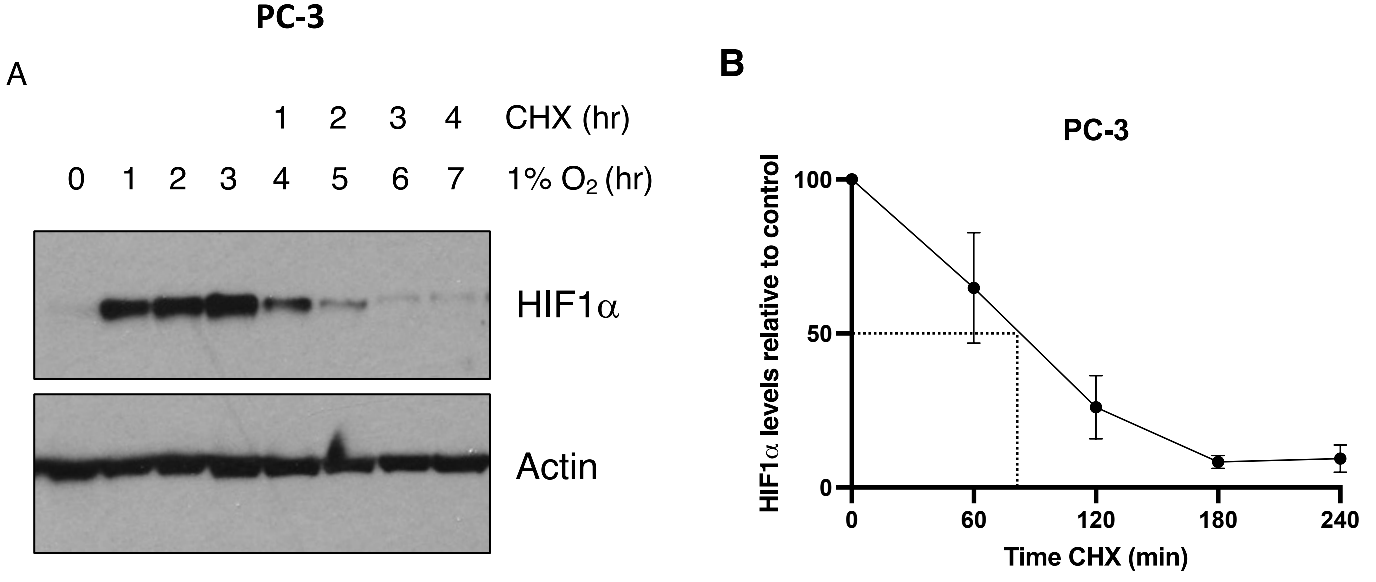

### Supplemental Figure 2

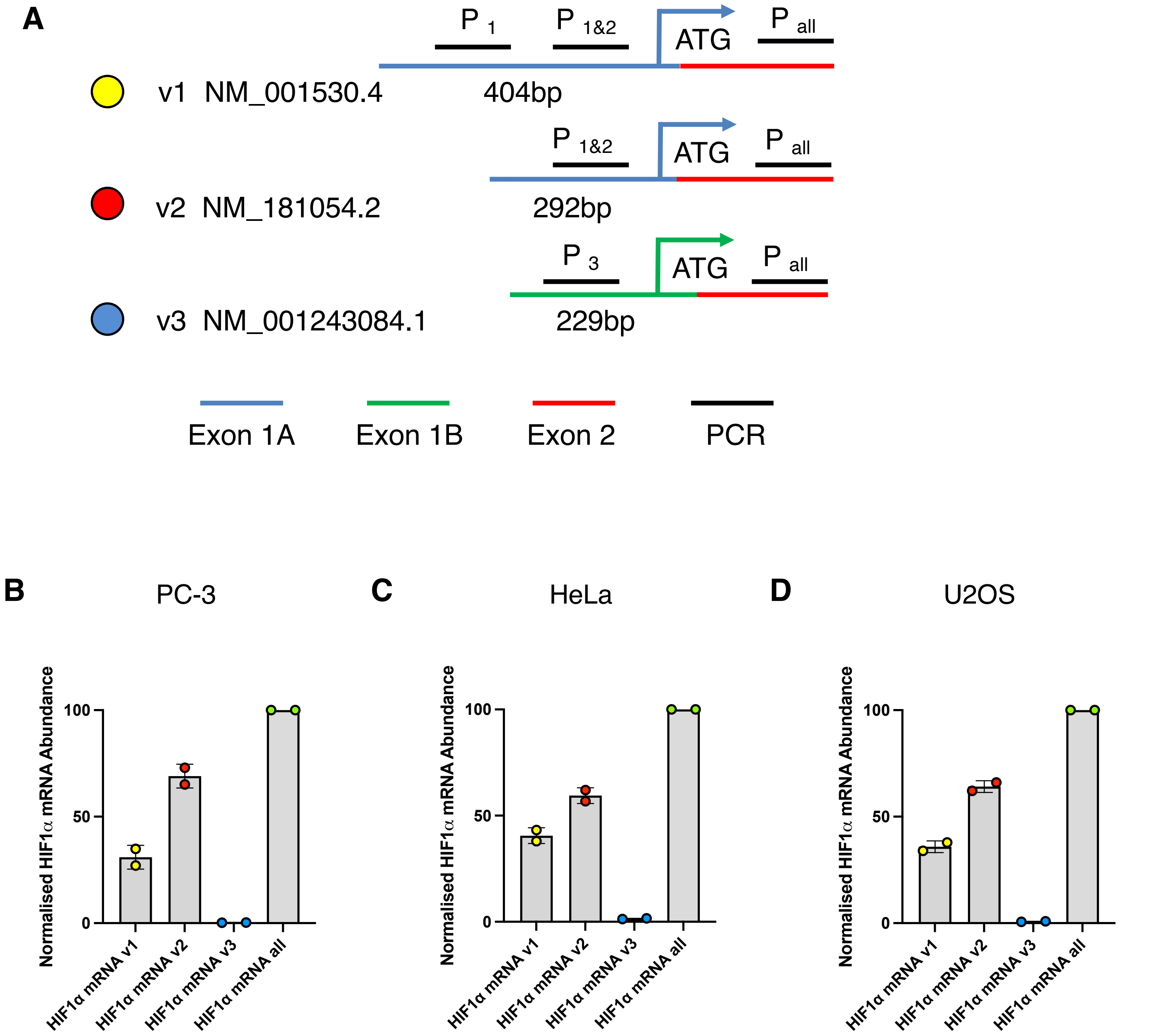

### Supplemental Figure 3

**A**

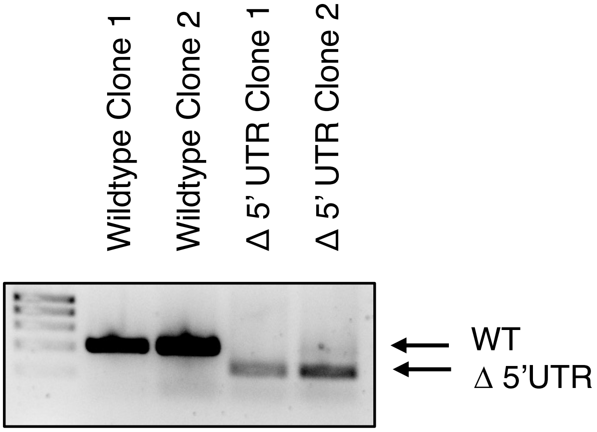


**B**

```
                                                     gRNA 1

Reference Seq -    1- CAGTGCTGCCTCGTCTGAGGGGAGAGGATCACCCTCTTCGTCGCTTCGGCCAGTGTGTCGGGCTGGGCCCTGACAAGCCACCTGAGGAGAGGCTCGGAGC -100

Control Cl. 1 -    1- CAGTGCTGCCTCGTCTGAGGGGAGAGGATCACCCTCTTCGTCGCTTCGGCCAGTGTGTCGGGCTGGGCCCTGACAAGCCACCTGAGGAGAGGCTCGGAGC -100

Control Cl. 2 -    1- CAGTGCTGCCTCGTCTGAGGGGAGAGGATCACCCTCTTCGTCGCTTCGGCCAGTGTGTCGGGCTGGGCCCTGACAAGCCACCTGAGGAGAGGCTCGGAGC -100

D 5'UTR Cl. 1 -    1- CAGTGCTGCCTCGTCTGAGGGGAGAGGATCACCCTCTTCGTCGC-------------------------------------------------------- -44

D 5'UTR Cl. 2 -    1- CAGTGCTGCCTCGTCTGAGGGGAGAGGATCACCCTCTTCGTCGC-------------------------------------------------------- -44



Reference Seq -  101- CGGGCCCGGACCCCGGCGATTGCCGCCCGCTTCTCTCTAGTCTCACGAGGGGTTTCCCGCCTCGCACCCCCACCTCTGGACTTGCCTTTCCTTCTCTTCT -200

Control Cl. 1 -  101- CGGGCCCGGACCCCGGCGATTGCCGCCCGCTTCTCTCTAGTCTCACGAGGGGTTTCCCGCCTCGCACCCCCACCTCTGGACTTGCCTTTCCTTCTCTTCT -200

Control Cl. 2 -  101- CGGGCCCGGACCCCGGCGATTGCCGCCCGCTTCTCTCTAGTCTCACGAGGGGTTTCCCGCCTCGCACCCCCACCTCTGGACTTGCCTTTCCTTCTCTTCT -200

D 5'UTR Cl. 1 -   44- ---------------------------------------------------------------------------------------------------- -44

D 5'UTR Cl. 2 -   44- ---------------------------------------------------------------------------------------------------- -44



                                             gRNA 2                                          Translation Start

Reference Seq -  201- CCGCGTGTGGAGGGAGCCAGCGCTTAGGCCGGAGCGAGCCTGGGGGCCGCCCGCCGTGAAGACATCGCGGGGACCGATTCACCATGGAGGGCGCCGGCGG -300

Control Cl. 1 —  201- CCGCGTGTGGAGGGAGCCAGCGCTTAGGCCGGAGCGAGCCTGGGGGCCGCCCGCCGTGAAGACATCGCGGGGACCGATTCACCATGGAGGGCGCCGGCGG -300

Control Cl. 2 —  201- CCGCGTGTGGAGGGAGCCAGCGCTTAGGCCGGAGCGAGCCTGGGGGCCGCCCGCCGTGAAGACATCGCGGGGACCGATTCACCATGGAGGGCGCCGGCGG -300

D 5'UTR Cl. 1 -   45- ---------------------------------------CTGGGGGCCGCCCGCCGTGAAGACATCGCGGGGACCGATTCACCATGGAGGGCGCCGGCGG -105

D 5'UTR Cl. 2 -   45- ---------------------------------------CTGGGGGCCGCCCGCCGTGAAGACATCGCGGGGACCGATTCACCATGGAGGGCGCCGGCGG -105
```

Supplemental Figure 4

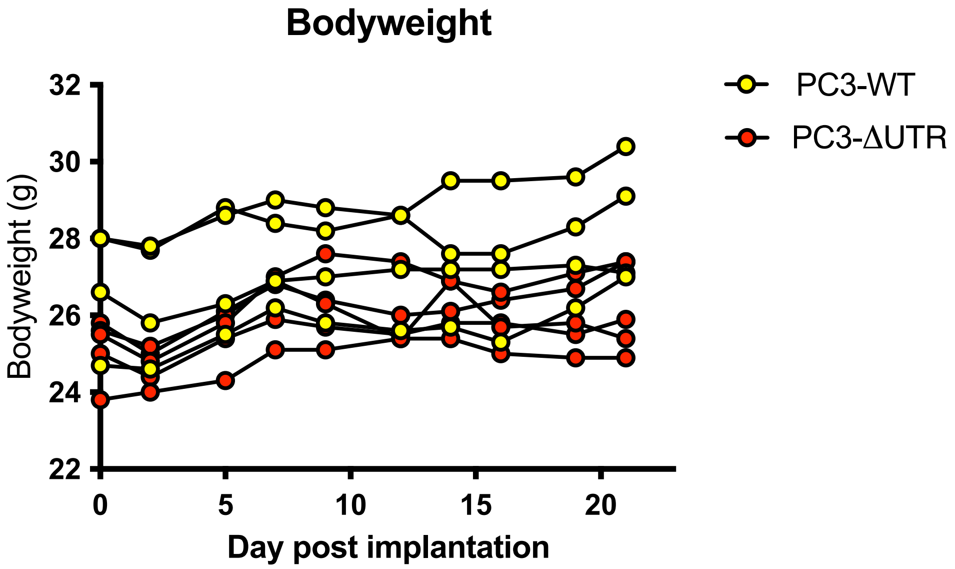

**Raw Blots for Figure 1B-D**

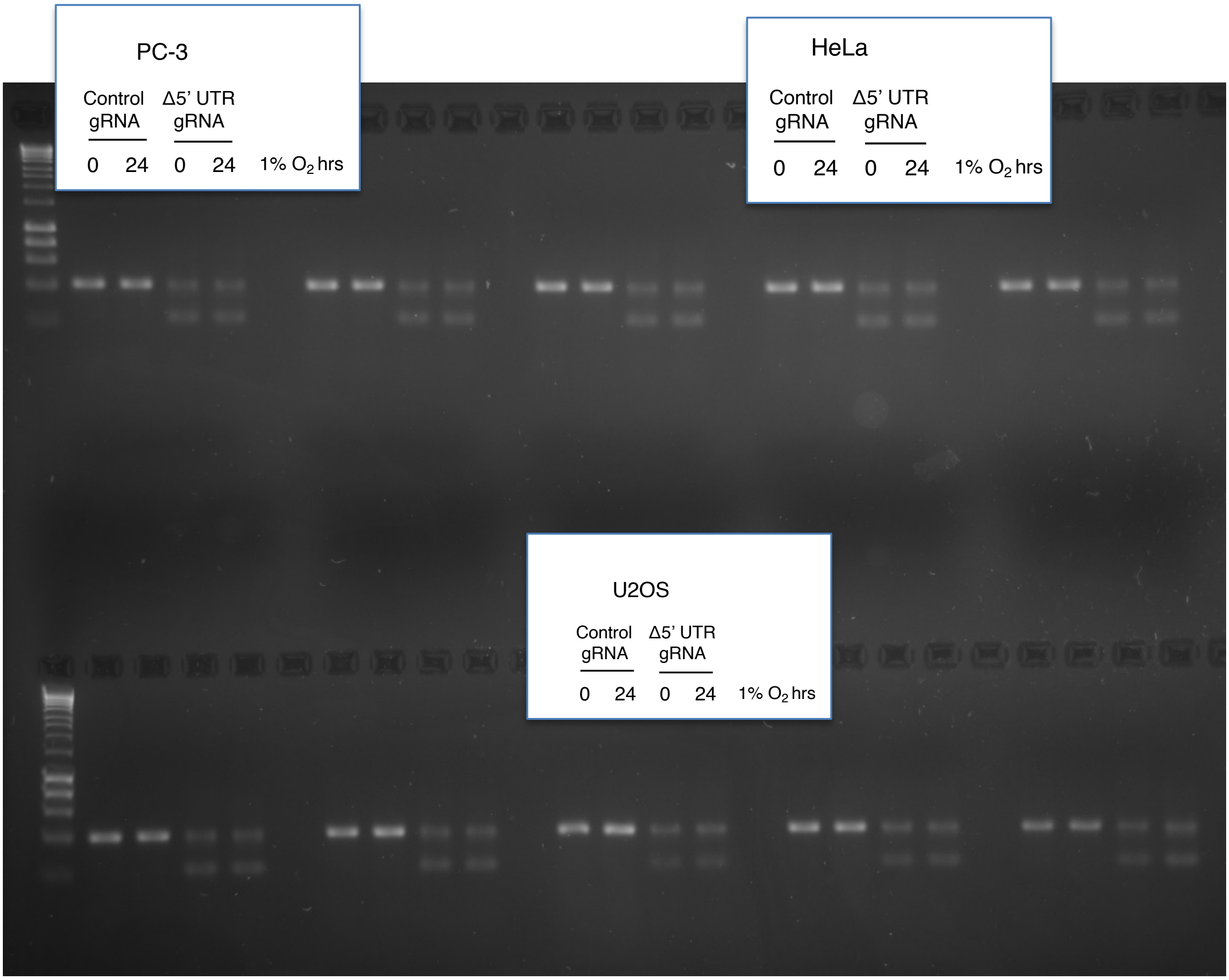

#### Full Blots for Figure 2A-C

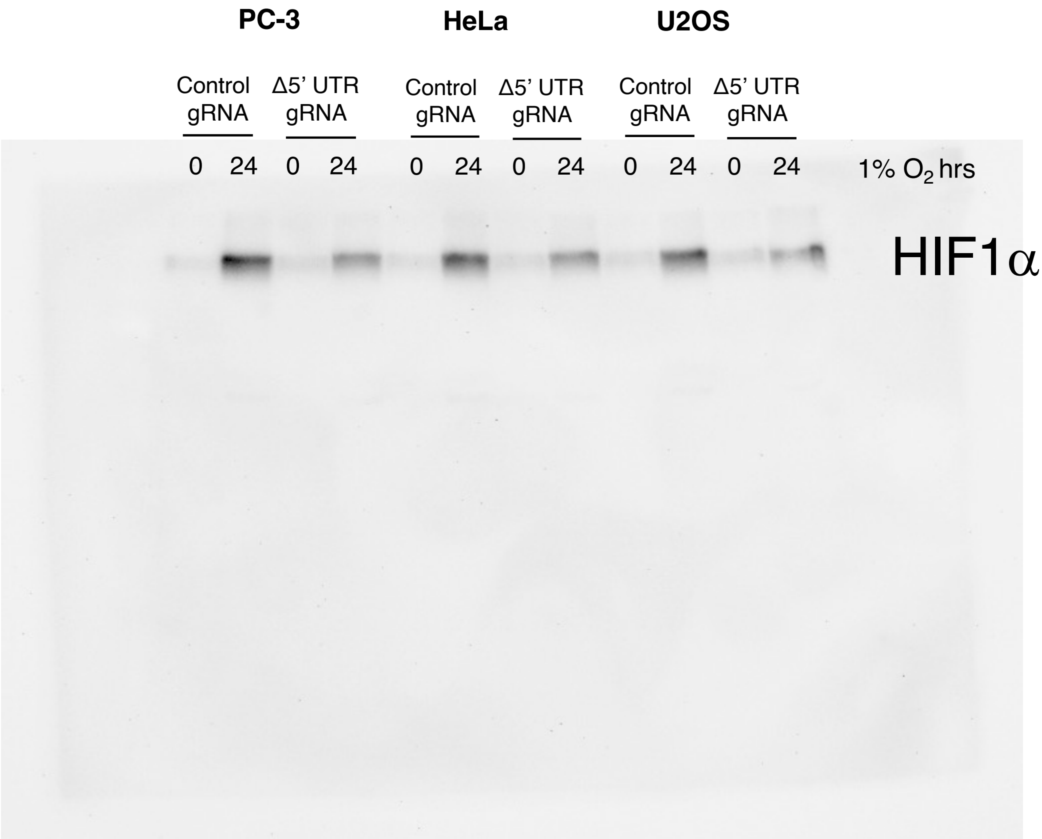


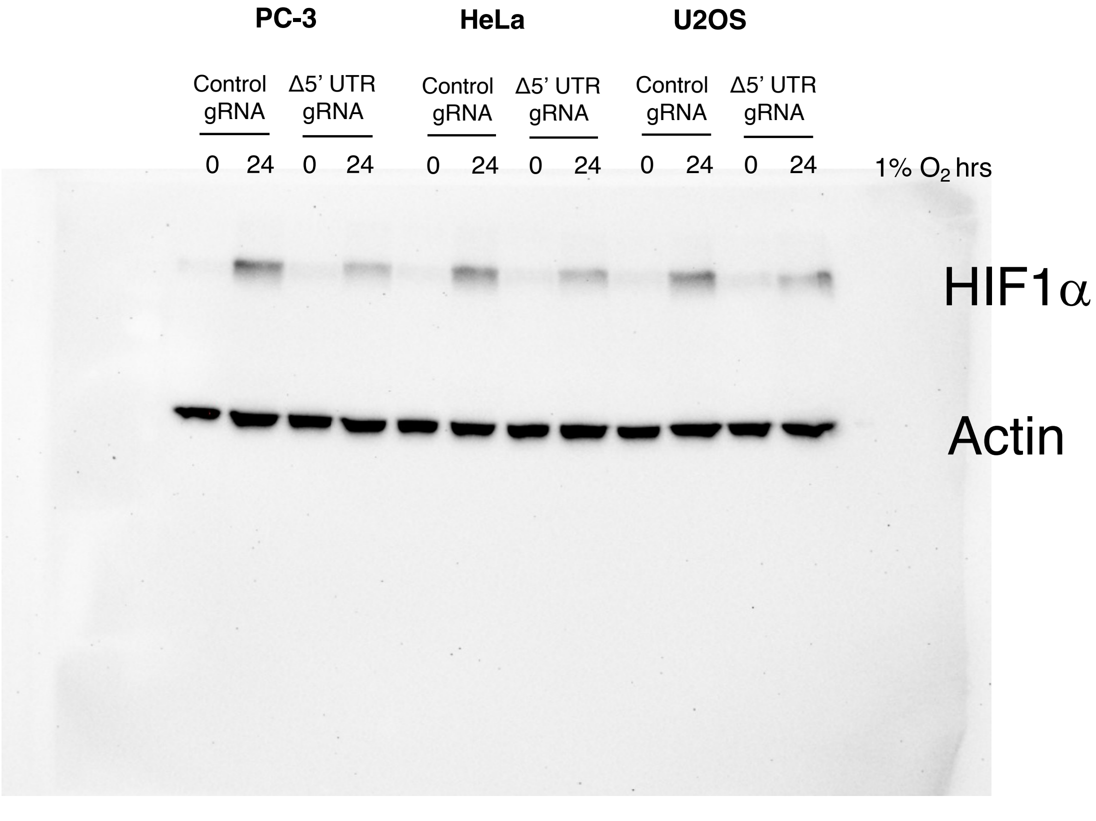

**Full Blots for Figure 3A**

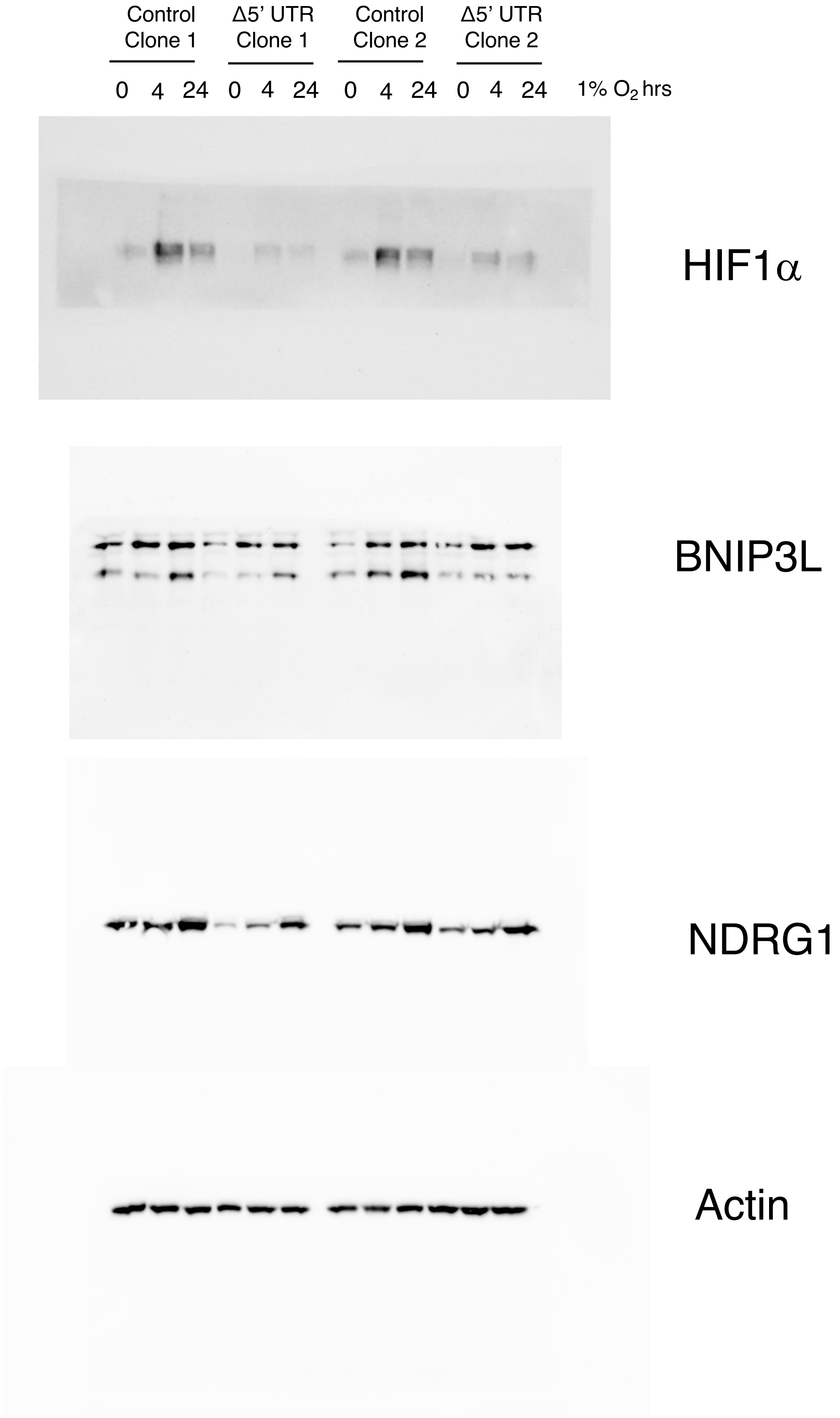

#### Full Blots for Figure 4C

Control Clone 1 | Δ5’ UTR Clone 1

0 24 0 24 1% $O_2$ hrs

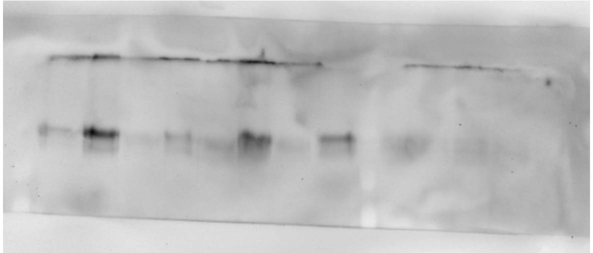

HIF1α

Control Clone 1 | Δ5’ UTR Clone 1

0 24 0 24 1% $O_2$ hrs

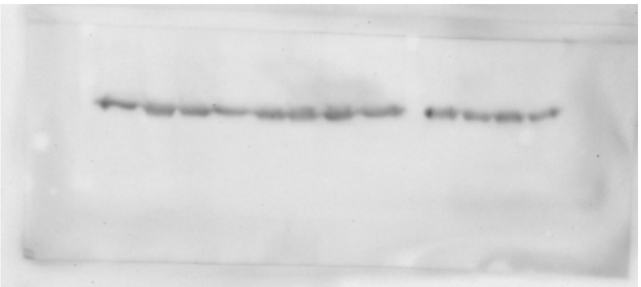

YB-1

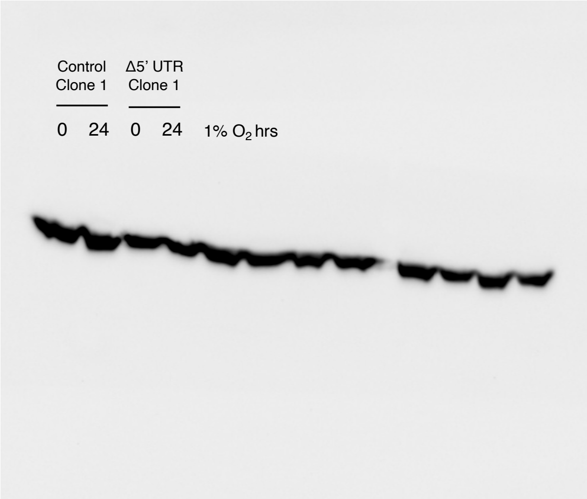


Actin

**Full Blots for Supplemental Figure 1**

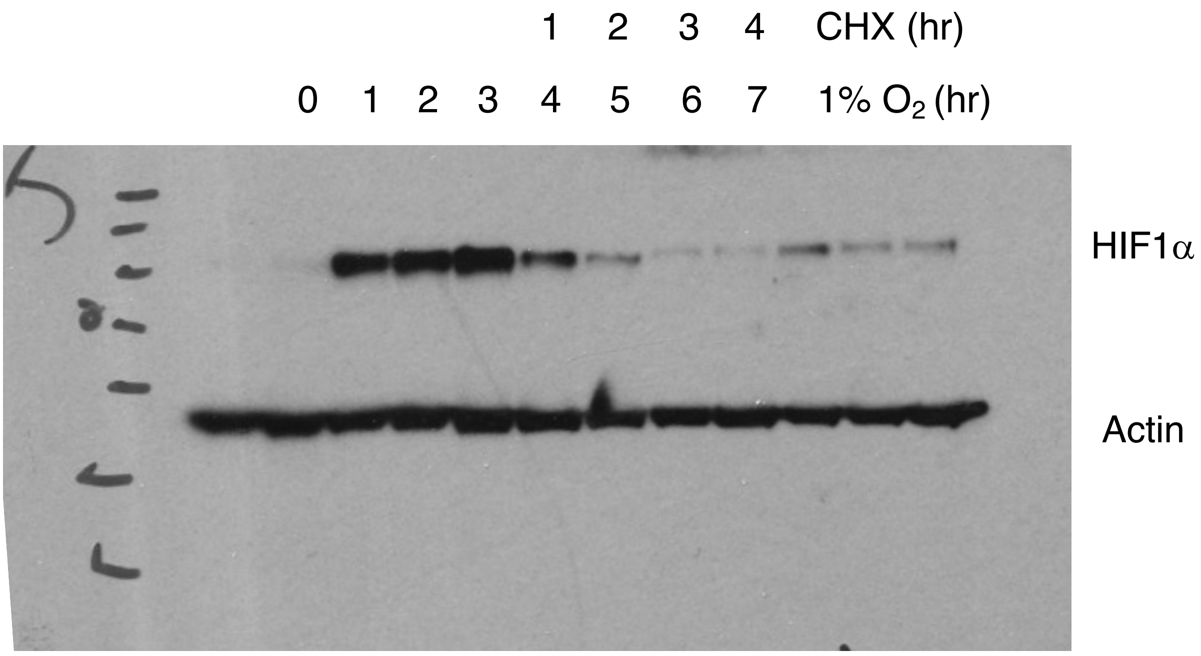

**Full Blots for Supplemental Figure 3**

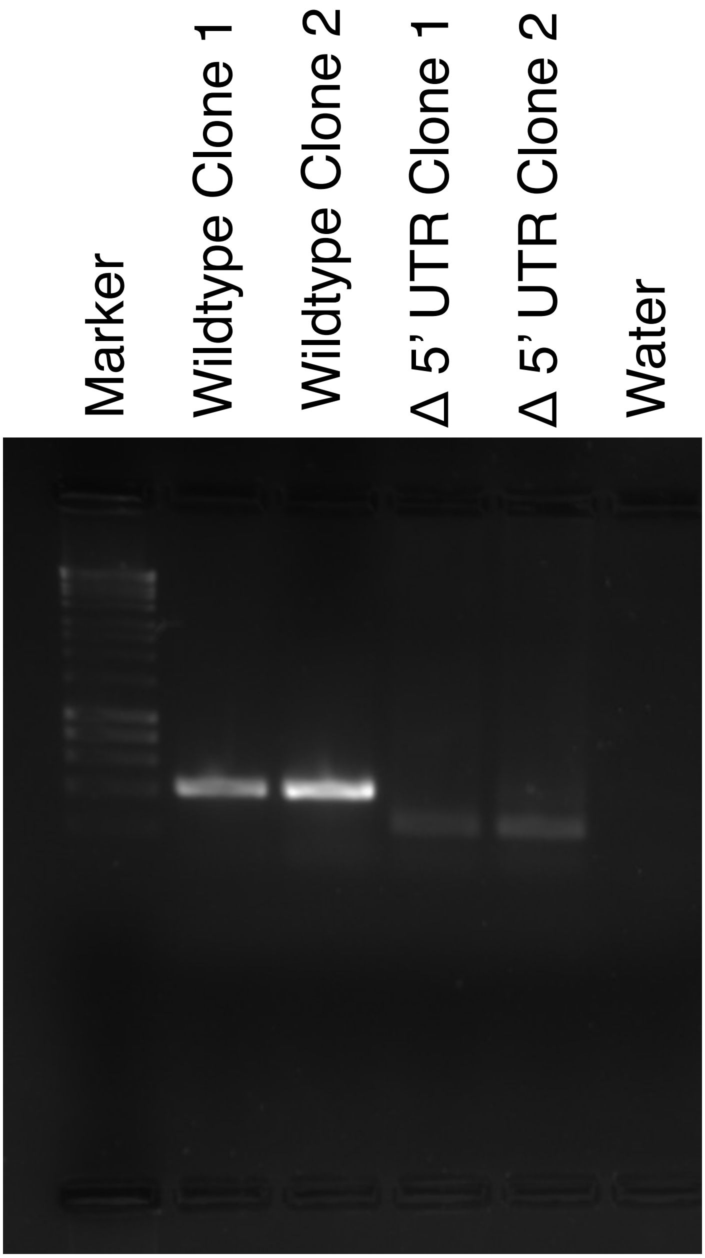

**Supplemental Figure 1**

(A) PC-3 cells were treated were exposed to 1% $O_2$ for the indicated times. DMSO or Cyclohexamide (100nM) was added after 3 hours hypoxia to block de novo protein synthesis. Whole-cell lysates (WCLs) prepared from these cells were subjected to immunoblot analysis to assess expression levels of the indicated proteins. (B) HIF1α levels were analysed by densitometry and normalised to b-actin levels. Relative HIF1α levels were plotted and the HIF1α half-life calculated.

**Supplemental Figure 2.**

**(A)** Schematic of the 3 HIF1α mRNA variants: NM_001530.4, NM_181054.2 and NM_001243084.1, designated v1, v2 and v3 in this paper. The lengths of the 5'UTRs are inidcated and the primer sets designed to investigate their relative levels are indicated. Primers P1 recognise v1, P1&3 recognise both v1 and v2; P3 recognise v3 and Ptot recognize exon2 which is common to all isoforms. Quantitative RT–PCR analysis of HIF1α mRNA isoforms from (B) PC-3, (C) HeLA and (D) U2OS cells.

**Supplemental Figure 3**

Analysis of monoclonal cell lines with deleted HIF1α 5' UTR. **(A)** PCR analysis of d UTR modified cell line demonstrating deletion of HIF1α UTR in all alleles **(B)** Sanger sequencing of D 5'UTR clones aligned to the wild type sequence showing the HIF1α 5; UTR wildtype sequence.

**Supplemental Figure 4**

WT PC-3 cells or 5'UTR PC-3 cells xenografts were implanted into nude mice. Weights of individual mice throughout the experiment were measured and plotted.

All raw image files for immunoblots and DNA gels.
